# The glycine locating at random coil of picornaviruses VP3 enhances viral pathogenicity by targeting p53 to promote apoptosis and autophagy

**DOI:** 10.1101/718130

**Authors:** Ruoqing Mao, Fan Yang, Dehui Sun, Xiaoli Zhou, Zixiang Zhu, Xuan Guo, Huanan Liu, Hong Tian, Keshan Zhang, Wen Dang, Qingfeng Wu, Xinwen Ma, Xiangtao Liu, Haixue Zheng

**Author notes:** Correspondence Author, Prof. Dr. Haixue Zheng. Lanzhou Veterinary Research Institute, Chinese Academy of Agricultural Science, No. 1, Xujiaping Road, Lanzhou, 730046, PR China.

## Abstract

Picornaviruses, comprising important and widespread pathogens of humans and animals, have evolved to control apoptosis and autophagy for their replication and spread. However, the underlying mechanism of the association between apoptosis/autophage and viral pathogenicity remains unclear. In the present study, VP3 of picornaviruses was demonstrated to induce apoptosis and autophagy. Foot-and-mouth disease virus (FMDV), which served as a research model here, can strongly induce both apoptosis and autophagy in the skin lesions. By directly interacting with p53, FMDV-VP3 facilitates its phosphorylation and translocation, resulting in Bcl-2 family-mediated apoptosis and LC3-dependent autophagy. The single residue Gly129 of FMDV-VP3 plays a crucial role in apoptosis and autophagy induction and the interaction with p53. Consistently, the comparison of rescued FMDV with mutated Gly129 and parental virus showed that the Gly129 is indispensable for viral replication and pathogenicity. More importantly, the Gly129 locates at a bend region of random coil structure, the mutation of Gly to Ala remarkably shrunk the volume of viral cavity. Coincidentally, the Gly is conserved in the similarly location of other picornaviruses, including poliovirus (PV), enterovirus 71 (EV71), coxsackievirus (CV) and seneca valley virus (SVA). This study demonstrates that picornaviruses induce apoptosis and autophagy to facilitate its pathogenicity and the Gly is functional site, providing novel insights into picornavirus biology.

## INTRODUCTION

Viral pathogenicity is frequently associated with the ability of the virus to kill host cells. Apoptosis, type I cell death and the main and most typical pattern of cell death, can be induced or inhibited by viral infection. In order to inhibit viral replication and dissemination, the host’s immune system immediately responds to viral invasion by initiating self-destructive apoptosis to curtail infection as an effective innate response. Viruses, however, have evolved a variety of strategies to regulate and control apoptosis in the host to ensure their continuous replication and release (1, 2). Apoptosis can be triggered by two main signaling pathways (intrinsic and extrinsic) regulated by various factors at multiple levels. The type II cell death is known as autophagic cell death (3). The relation between autophagy and virus is intricate. As an antiviral mechanism, autophagy proteins can influence innate and adaptive immune response, resulting in autophagy-mediated viral degradation and ultimately inhibit viral replication and release. Also, in some cases autophagy as a cellar survival mechanism limits virus-induced apoptosis, to protect neighboring uninfected cells (4, 5). In contrast to antiviral function, the autophagosome with cellular membranous structure as a platform for the replication and translation of viral membrane-associated replication factories, can promote viral replication (6–8).

The relationship between apoptosis and autophagy is complicated, and in some scenarios they crosstalk with eachother via several molecular nodes, such as target of rapamycin (TOR), Beclin 1, caspase, Flice inhibitory protein (FLIP), death-associated protein kinase (DAPK) and p53 (9, 10). The p53 tumor suppressor is considered a crucial mediator of apoptosis, autophagy, cell cycle, metabolism and senescence in response to stimulating stresses (11, 12). It can be activated by internal and external stimuli that promotes its accumulation in a stable and activated form by phosphorylation, acetylation or SUMOylation. The stabilized p53 in turn regulates many pro-apoptotic genes such as Bax, Bad, Bid, Fas, and PUMA (13). In recent years, several reports have focused on the relationship between p53 and autophagy. p53 can target DRAM and Isg20L1 to activate autophagy or enhance mTOR activity to inhibit autophagy (14). Conversely, autophagy represses p53 by suppressing oxidative stress or preventing DNA damage.

Picornaviruses comprise a large number of non-enveloped small RNA viruses, including hepatitis A virus (HAV), poliovirus (PV), foot-and-mouth disease virus (FMDV), enterovirus 71 (EV71), coxsackievirus (CV) and seneca valley virus (SVA), which are important pathogens of humans and animals (15). The single positive-stranded RNA genome of picornaviruses consists of a single open reading frame (ORF) encoding a polyprotein that is post-translationally processed into four structural proteins (VP1, VP2, VP3 and VP4) and eight non-structural proteins (Lpro, 2A, 2B, 2C, 3A, 3B, 3C, and 3Dpol) (15). As other viruses, picornaviruses have evolved to control apoptosis for viral replication and spread (16, 17). Coxsackievirus 2A protease can cleave DAP5 to enhance viral replication and apoptosis (18). The PV 3A protein inhibits TNF-induced apoptosis (19). 3C can induce apoptosis in PV-infected cells by caspase activation (20). EV71 2B enhances apoptosis by inducing conformational activation of BAX (21). 3C enhances apoptosis by Pinx1 cleavage (22).

Autophagy also plays important roles in picornaviruses infection or replication. EV71 induced autophagy promotes viral replication (23), and the promyelocytic leukemia (PML) represses EV71 replication by inhibiting autophagy (24). PV proteins 2BC and 3A are contributing to the modification of LC3 (25).

The FMDV is another well-known etiological agent belonging to the genus Aphthovirus in the Picornaviridae family; its associated disease is notorious for colossal and disastrous impacts on livestock characterized by fever, lameness and vesicular lesions (26). The FMDV causes cytopathic effects (CPE) in infected host cells, with dramatic structural and morphological changes, which underlie viral pathogenicity without clearing mechanisms (26, 27). Concomitant with CPE, cell death is commonly observed in infected cells. There are only two proteins, VP1 and 2C, which have been proven to play important roles in FMDV-induced cellular apoptosis (28–30). VP2 interacts with HSPB1 to induce autophagy and enhance viral replication (31). However, how host apoptosis and autophagy affect viral pathogenicity is unclear. In this study, we firstly identified and demonstrated the structural protein VP3 of PV, FMDV and SVA as novel proteins inducing apoptosis and autophagy. To understand the further mechanism, taking FMDV as a model and FMDV-VP3 was shown to directly interact with p53, facilitating p53 phosphorylation, translocation into the mitochondria and interaction with Bad, which result in cytochrome c-mediated apoptosis and LC3-dependent autophagy. Furthermore, the single residue Gly129 of VP3 was essential for the VP3- and FMDV-induced apoptosis and autophagy, as well as the interaction between VP3 and p53. Meanwhile, apoptosis and autophagy were shown to be a critical mechanism promoting FMDV replication and pathogenesis. More importantly, three dimensional (3D) structure models showed that the mutation of Gly to Ala remarkably shrunk the volume of viral cavity. Coincidentally, at the similar location of other picornaviruses, including CV, EV71, SVA and PV, Gly is conservative. All data indicate that the ability of cell death induction may be conserved amongst picornaviruses and this may depend on the conservative structure of VP3.

## MATERIALS AND METHODS

### Cells and viruses

The hTERT-BTY cell line was established from primary bovine thyroid (BTY) cells in our laboratory (Invention Patent, China, ZL201410421962.4. CCTCC, C2014109) (32). The cells were cultured in Dulbecco’s Modified Eagle Medium/Nutrient Mixture F-12 (DMEM/F12; Gibco) supplemented with 10% fetal bovine serum (Gibco), 10 μg/mL insulin, 100 U/mL penicillin and 100 μg/mL streptomycin (Sigma). The baby hamster kidney (BHK-21, ATCC, CCL-10), porcine kidney epithelial (PK-15, ATCC, CCL-33) and human embryonic kidney 293T (HEK293T, ATCC, CRL-3216) cell lines were maintained in minimum essential medium (MEM) supplemented with 10% fetal bovine serum (Gibco) and 100 U/mL of penicillin-streptomycin. All cells were cultured at 37°C in a humid environment with 5% CO_2_. The mycoplasma test was performed to ensure that the cell lines were mycoplasma-free.

Type O FMDV strain O/BY/CHA/2010 (GenBank: JN998085.1) was obtained from the Chinese National Reference Laboratory for Foot and Mouth Diseases, and propagated in BHK-21 cells. Viral infection was performed according to the standard procedure(33). Then, hTERT-BTY cells at 90% confluence were infected with FMDV; after 1h adsorption at 37 °C, the cells were washed twice with PBS and cultured continuously in MEM without FBS at 37 °C and 5% CO_2_.

### Reagents, plasmids and antibodies

Cell apoptosis was analyzed with an Annexin V-FITC/propidium iodide (AnnV/PI) apoptosis assay kit (Invitrogen, Carlsbad, CA). Mitochondrial membrane potential was analyzed with a Mitoprobe™ JC-1 Assay kit (Invitrogen). Nuclear condensation was assessed by staining with the Hoechst^®^ dye (Invitrogen). The full-length cDNAs of type O FMDV strain proteins were respectively cloned into the pCAGGS-Flag vector to express viral proteins in eukaryotic expression plasmids. A series of truncated or mutated VP3 plasmids were generated by site-directed mutagenesis PCR. The VP3 of PV (GenBank: FJ769385.1) and SVA (GenBank: KY747510.1) were respectively cloned into the pCMV-Flag vector to express viral proteins in eukaryotic expression plasmids. The human caspases 3, 7 and 8, and JNK, AKT, P53, Bad, Bid, Bax, Bcl-2 and XIAP were cloned into pCDNA 3.1-Myc or pCMV-HA. Antibodies specific for β-actin (sc-47778), caspase-3 (sc-1225), 8 (sc-6139), caspase-9 (sc-7885), Bax (sc-493), Bcl-2 (sc-492), Cyto-c (sc-7159), AKT (sc-8312), JNK (sc-571), Bad (SC-8044, immunofluorescence), p53 (sc-99) and p-p53 (sc-51690) were purchased from Santa Cruz Biotechnology (Santa Cruz, CA, USA). Antibodies specific for Cox IV (ab14744), Bad (ab90435), p53 (ab61241, immunofluorescence), mouse or rabbit antibodies specific for HA (ab1424), Flag (ab1162) and Myc (ab32) were purchased from Abcam (Abcam, Cambridge, UK). Antibodies specific for LC3 (PM036) were purchased from MBL (MBL, JP). Polyclonal antibodies specific for O type FMDV and monoclonal antibodies targeting O type FMDV-VP3 were prepared by our laboratory (unpublished data).

### Apoptosis assay, mitochondrial membrane potential detection and measurement of nuclear condensation

Early and late stage cell apoptotic events were analyzed by AnnV/PI staining. The externalized phospholipid phosphatidylserine, a typical marker of cells undergoing apoptosis, is stained by Annexin V-FITC, whereas PI binds the DNA of late apoptotic and necrosis cells.

Cells were seeded in 6-well plates and cultured to 90% confluence. After viral infection or transfection, cells were detached by trypsin without EDTA at different time points. All cells including those in the supernatant were collected and resuspended in binding buffer at a density of 1×10^6^ cells/mL. The cell suspensions were stained with Annexin V-FITC and PI at 4°C in the dark, and fluorescence was measured by flow cytometry. Ten thousand cells in each sample were analyzed.

The disruption of active mitochondria, causing membrane potential changes and alterations of the oxidation-reduction potential, is a distinctive characteristic of early apoptosis. The membrane-permeant JC-1 dye is commonly used in apoptosis assays. After viral infection or plasmid transfection, the cells were stained with 2μM JC-1 for 15 min at 37°C, 5% CO_2_. Then, the cells were washed with PBS and analyzed by flow cytometry with excitation at 488 nm and emission at 530 nm and 585 nm, respectively.

Hoechst dye is often used to observe condensed pycnotic nuclei in apoptotic cells. After plasmid transfection or viral infection, the cells were fixed with 4% paraformaldehyde, incubated with Hoechst 33342 staining solution for 5-10 minutes, washed three times with PBS, and subjected to analysis by fluorescence microscopy.

### Western blot, immunofluorescence and Co-immunoprecipitation

Total protein from cells was extracted with cell lysis buffer for Western blot and IP (Beyotime, Shanghai, China). Equal amounts of total protein were resolved by SDS-PAGE and transferred onto a PVDF membrane (Millipore). The membrane was then blocked with horse blocking buffer (bioWORLD, USA) and sequentially incubated with specific primary and secondary antibodies. Enhanced chemiluminescence detection reagents (Thermo) were used to visualize target proteins.

For immunofluorescence, cells were grown on confocal dishes and transfected with various plasmids. The Mitochondrion-selective probe (GeneCopoeia, Rockville, USA) was used to stain cell mitochondria according to the manufacturer’s instructions. After staining, the cells were fixed with 4% paraformaldehyde (Sigma) for 1 h. After three PBS washes, the cells were permeabilized with 0.1% Triton-100 (Sigma) for 20 min at room temperature and blocked with 5% BSA for 1 h at 37°C. The specimens were next incubated with primary antibodies followed by fluorochrome-conjugated secondary antibodies. Finally, the cells were incubated with DAPI-Fluoromount-G (Solarbio, Beijing, China) and visualized under a confocal laser scanning microscope (TSC SP5 Leica).

Co-immunoprecipitation assays were performed as described previously (34). Briefly, the cells were co-transfected with the indicated plasmids. Then, they were lysed and incubated with various monoclonal antibody-conjugated agarose beads, respectively, overnight at 4°C on a rotary vibrator. The beads were washed with lysis buffer three times and boiled. Proteins in the supernatants were analyzed by Western blot.

### Real-time PCR

One step quantitative real-time RT-PCR (rRT-PCR) was performed to detect viral RNA as described previously.(35, 36) A conserved region in the FMDV 3D gene was chosen for primer and TaqMan probe design. Total RNA was extracted with TRIzol Reagent (Invitrogen) and One Step Primescript TM RT-PCR Kit was used to determine FMDV copies. The experiments were performed at least three times, and a threshold cycle (CT) value was assigned to each PCR reaction. Then, standard curves showing a linear relationship between CT and FMDV copies were established to quantify FMDV.

### Construction of apoptotic site-mutated recombinant viruses

Based on the reverse-genetics system established in our laboratory (37, 38), we constructed an infectious cDNA of O/CHA/99, termed pO-FMDV. The P1 coding sequence of O/BY/CHA/2010 was selected as the site for exchange with the corresponding region of pO-FMDV using the restriction endonucleases AfIII and Clal (New England Biolabs, Ipswich, Massachusetts, USA); the recombinant plasmid was named prVP3/FMDV. Site-directed mutagenesis performed with specific primers (VP3-129-F, 5’-GCCCCTCCGGCTATGGAGCCGCCCAAAACAC-3’; VP3-129-R, 5’-CGGCTCCATACGCGGAGGGGCATACGCAATC-3’) was used to mutate VP3 Gly129 to Ala. The positive plasmid with the VP3 G129A substitution was digested with AfIII and Clal, and reintroduced into the pO-FMDV vector, producing the recombinant plasmid PrVP3-129/FMDV. Sequencing and purification of the above resulting plasmids were performed as described previously (33).

The purified plasmids were extracted with NucleoBond^®^ Xtra Maxi Kit (Macherey-Nagel, Düren, Germany), and transfected into BHK-21 cells with Lipofectamine™ 2000 (Invitrogen, Carlsbad, CA) following the manufacturer’s protocol. The supernatants were harvested after 48 h and subjected to three freeze-thaw cycles. Then, the viruses recovered from the abovementioned supernatants were harvested by centrifugation at 5,000×g for 10 min at 4°C, and passaged 30 times in BHK-21 cells. The recombinant viruses were stored at −80°C for future use.

### Virus internalization

hTERT-BTY and PK-15 cells (5×10^5^ cells/well) were infected with the two rescued viruses at a dose of 1×10^8^ viral genomic RNA copies for 1 h at 37°C, and unabsorbed viruses were washed with ice-cold Hanks balanced salt solution (HBSS) three times. To remove surface-bound viruses, the cells were digested with trypsin-EDTA (GIBCO) for 5 min, and washed three times with HBSS.

### Virus titration and plaque assay

The plaque assay was performed as described previously.(35) Cells were seeded in 6-well plates and cultured to 90% confluence. Then, 10-fold dilutions of viruses were inoculated into the cells. The medium was removed after 1 h of adsorption, and cells were overlaid with 50% 2×MEM supplemented with 2% FBS and 50% Tragacanth. The cells were incubated at 37°C for 48 h, fixed with methanol and acetone (1:1), and stained with crystal violet (Sigma).

### Hydrophobicity prediction and homology modeling

The ExPASY-ProtScale tool (https://web.expasy.org/protscale/) was used to predicate the Kyte & Doolittle hydrophobicity of viral proteins. The SWISS MODEL (http://swissmodel.expasy.org/) was used for 3D protein structure predicting. 3D structure of the mutated VP3 was generated by PyMOL software.

### Guinea pig challenge experiments

Animal experiments were performed at the Biosafety Level 3 laboratory of LVRI, Chinese Academy of Agricultural Sciences (Permission number: SYXK-GAN-2004-0005). All animal experiments were approved by the Gansu Animal Experiments Inspectorate and the Gansu Ethical Review Committee (License no. SYXK [GAN] 2010-003). Animals in this study were humanly treated and euthanized by injection of sodium pentobarbital at the end of the experiments. Female Hartley guinea pigs, weighing 300-350 g and serologically negative for FMDV, were obtained from Lanzhou Veterinary Research Institute (China). The titers of the two rescued viruses were adjusted to 8.0 TCID50/mL. According to the principle of randomization, guinea pigs were divided into 16 groups (n=5 per group; this sample size meets the basic requirements for statistical analysis). Groups 1-8 were challenged intradermally and subcutaneously by footpad injection of a series of 10-fold dilutions from 0 (200μL undiluted virus/animal at a titer of 8.0 TCID50/mL) to −8 (200μL 10^8^-fold diluted virus/animal) of rVP3-129/FMDV per guinea pig. Groups 9-16 were challenged with rVP3/FMDV at the same dosage by the same procedure. All animals were assessed daily for signs of illness, and clinical signs were scored as follows by the double-blind method: no local red swelling and heat, 0; red swelling at the original injection site of one footpad, 1; red swelling at the original injection site of both footpads, 2; vesicles at one footpad, 3; red swelling at the original injection site of one footpad and vesicles at the other footpad, 4; vesicles at both footpads, 5.

Heparinized blood samples were collected on days 0, 3, 6, 10 and 15 post-challenge, respectively, and serum antibodies against FMDV were measured with a guinea pig FMDV IgG ELISA kit (Jianglaibio, China, Shanghai) according to the manufacturer’s instructions.

### Hematoxylin-eosin staining and the TUNEL-assay

Guinea pigs were euthanized by intravenous injection of sodium pentobarbital. Then, postmortem examination was performed, and tissue samples (heart, liver, spleen, lung, kidney, mesenteric lymph nodes, submaxillary lymph nodes and pathologic tissues) were collected. All samples were fixed with 4% paraformaldehyde for at least 24 h, dehydrated, and paraffin embedded. The treated tissues were sectioned at 4μm and stained with hematoxylin and eosin (H&E).

To detect DNA fragmentation by labeling the 3’ – hydroxyl termini in the double-strand DNA breaks generated during apoptosis, the TUNEL assay was performed with TUNEL Apoptosis Assay Kit (Roche). Briefly, slides were deparaffinized, rehydrated, and incubated with 20μg/mL proteinase K for 25 min at 37°C for protein digestion. Then, the labeling mixture was added to the sections and incubated at 37°C for 2 h, and sections were rinsed with PBS. After nuclear staining, the sections were covered with mounting medium. Finally, images were acquired on an imaging system (Pannoramic MIDI/P250, Hungary), and the percentage of apoptotic cells was determined with Image pro plus 6.0.

### Tissue immunohistochemistry and immunofluorescence

Paraffin embedded tissues were sectioned at 2-3μm. Sections were dewaxed and pretreated as previously described (39). The blocked tissues were incubated with primary antibodies. For immunohistochemistry, the slides were incubated with biotinylated secondary antibodies, followed by DAB (ZSBIO, China) staining. Then, the slides were counterstained with Mayer’s hematoxylin (ZSBIO, China). For immunofluorescence, the slides were incubated with fluorophore-conjugated secondary antibodies, and cell nuclei were stained with DAPI. Images were acquired under a BA200 Digital microscope (Motic, China) or a confocal laser scanning microscope. Pearson’s correlation analysis was performed with Image pro plus 6.0.

### Statistical analysis

Student’s t-test was performed using the SPSS 7.0 software. The level of significance is shown in figures.

## RESULTS

### Predicting of FMDV, PV and SVA proteins identifies VP3 as inducer of apoptosis and autophagy

The function of proteins to regulate cell death is often related to its hydrophobic regions. The ExPASY-ProtScale tool was used to predicate the hydrophobic features of FMDV, PV and SVA proteins. By synthesizing high hydropathicity score and number of amino acids with the top score, VP3 of above three viruses were chosen for further study (Figure 1A). Three dimensional structures of FMDV-VP, PV-VP3 and SVA-VP3 were predicated using SWISS-MODEL tool. The models show that the three VP3 proteins have similar structure (Figure 1B). To explore whether the above proteins regulate host cell death, the expression levels of cleaved caspase 3 and LC3 were examined in VP3 proteins-transfected HEK293T cells. The results showed that all three VP3 significantly promoted activation of caspase 3 (Figure 1C) and upregulated LC3-II expression (Figure 1E). VP3-induced apoptosis was also verified by Annexin V-FITC/propidium iodide (AnnV/PI) staining and flow cytometry (Figure 1D). These results show that VP3 of FMDV, PV and SVA can induce both apoptosis (type I cell death) and autophagy (type II cell death).

**Figure 1.**
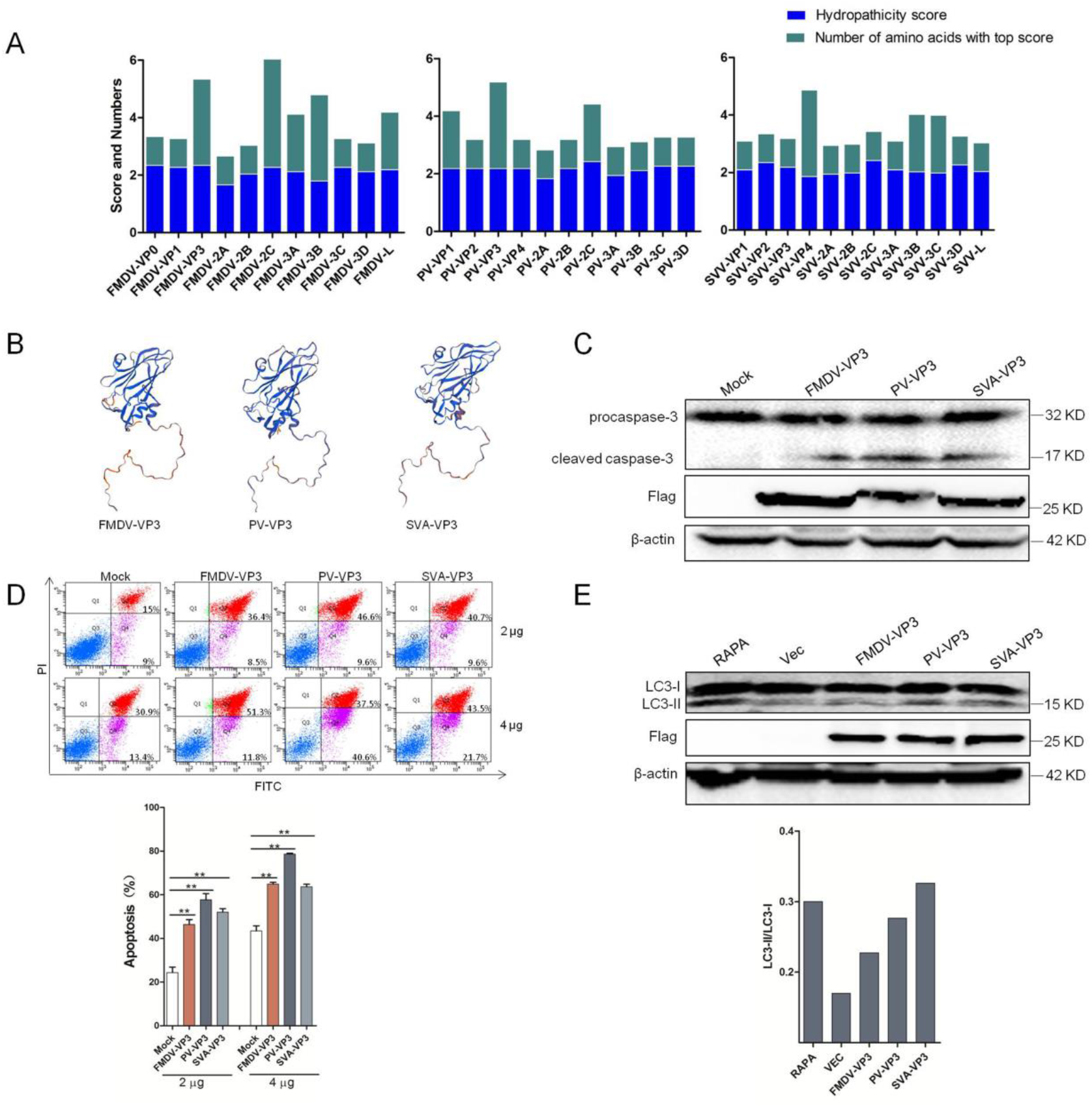
Prediction and identification of viral cell death regulation proteins. (A) Kyte & Doolittle hydrophobicity prediction of FMDV, PV and SVV proteins by ProtScale analysis at Expasy. (B) Three dimensional structures of FMDV-VP3 (PDB ID: 4IV1),PV-VP3 (PDB ID: 5KUO) and SVA-VP3 (PDB ID: 6ADS) were predicted using SWISS-MODEL tool. (C) HEK-293T cells (10^5^ cells/well) were transfected with empty vector or FMDV-, PV- and SVA-VP3. At 48 h.p.t., the expression levels of procaspase-3, cleaved caspase-3 and Flag-VP3 were detected by Western blot. (D) HEK-293T cells (10 cells/well) were transfected with increasing quantities of empty vector or FMDV-, PV- and SVA-VP3 (2μg or 4μg). Apoptosis was detected by Annexin V-FITC/PI staining and FCM at 48 h.p.t.. (E) HEK-293T cells (10^5^ cells/well) were transfected with empty vector or FMDV-, PV- and SVA-VP3. At 48 h.p.t., the expression levels of LC3-I, LC3-II and Flag-VP3 were detected by Western blot and the gray intensities of LC3-II/LC3-I were analyzed. Data are mean ± SD (n=3). Statistical significance was analyzed by Student’s t-test: *P<0.05, **P<0.01.

### FMDV infection induces apoptosis and autophagy in vivo

Cell death is commonly observed in FMDV-infected cells in vitro and in vivo. To confirm the type of FMDV-induced cell death, immunofluorescent staining on footpad lesions of FMDV-infected guinea pigs was performed. FMDV infection significantly upregulated the expression of LC3I/II and cleaved-caspase3 (Figure 2A). The images were further analyzed by HISTOQUEST software. The results showed that the percentage of LC3-I/II- and cleaved-caspase3-positive cells, whether in epidermis or dermis, was considerably higher in FMDV-infected guinea pig than PBS group. And the level of cleaved-caspase3 upregulation is more significant expressive than LC3I/II (Figure 2B). All these results indicate that FMDV can induce both apoptosis and autophagy in vivo, but the main type of cell death is apoptosis.

**Figure 2.**
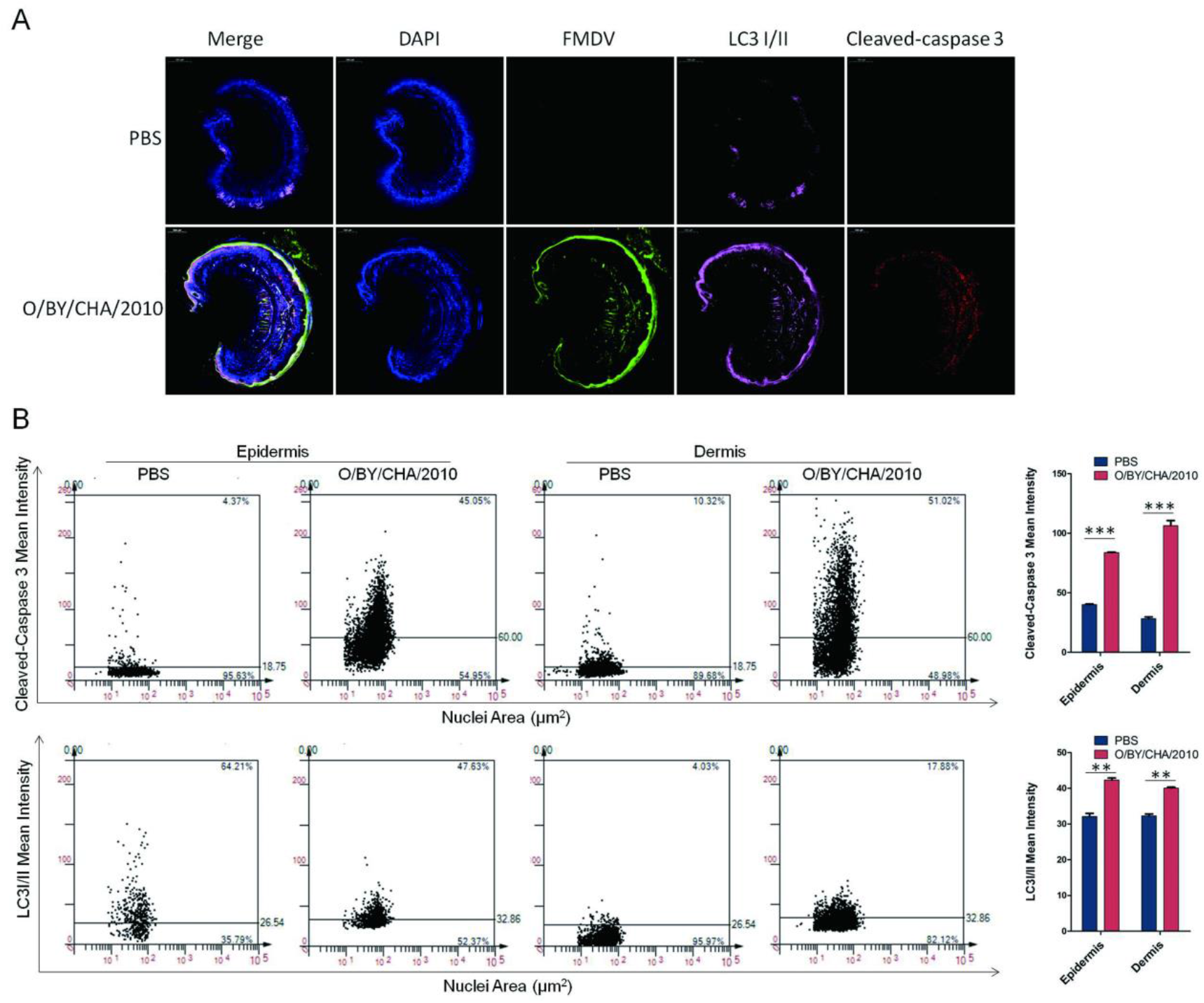
FMDV induces apoptosis and autophagy. (A) Guinea pigs were challenged with FMDV (O/BY/CHA/201 0, 8.0 TCID50/mL, 200μL/animal) or PBS. At 7 d.p.c., the lesions were incubated with the indicated antibodies and examined by confocal microscopy. DAPI, blue; FMDV, green; cleaved-caspase3, red; LC3-I/II, pink. (B) The cleaved-caspase3 and LC3-I/II positive cells of epidermis and dermis were analyzied respectively by HISTOQUEST soteware. Data are mean ± SD (n=3). Statistical significance was analyzed by Student’s t-test:*P<0.05, **P<0.01, ***P<0.001; d.p.c., days post challenge.

### Verified VP3 protein of FMDV as a strong inducer of cellular apoptosis

For the reason that apoptosis is the mian cell death type of FMDV-infected cells, viral apoptosis mechanisms were firstly studied. To investigate whether the FMDV could induce apoptosis in host cells in vitro, bovine thyroid (hTERT-BTY) cells were infected with O/BY/CHA/2010, and the FMDV caused clear CPE observed by light microscopy at 24 h post-infection (h.p.i.) as well as significant apoptosis analyzed by Annexin V-FITC/propidium iodide (AnnV/PI) staining and flow cytometry (Figure 3A). Next, all the FMDV proteins were screened for their abilities to induce apoptosis. Western blotting confirmed the successful expression of FMDV proteins (Figure 3B); among them, VP1, VP3 and 3A promoted apoptosis in hTERT-BTY cells at 24 and 48 h post-transfection (h.p.t.) (Figure 3C). The same phenomenon was observed in the porcine kidney epithelial PK-15 cell line. However, the baby hamster kidney BHK-21 cell line, a well-established cell line for FMDV propagation, was not an ideal model for apoptosis study due to its sensitivity to transfection; indeed, transfection of the vector also led to significant cellular apoptosis. Further, multifaceted approaches were used to evaluate the apoptosis-inducing activity of FMDV-VP3 (Figure S1).

**Figure 3.**
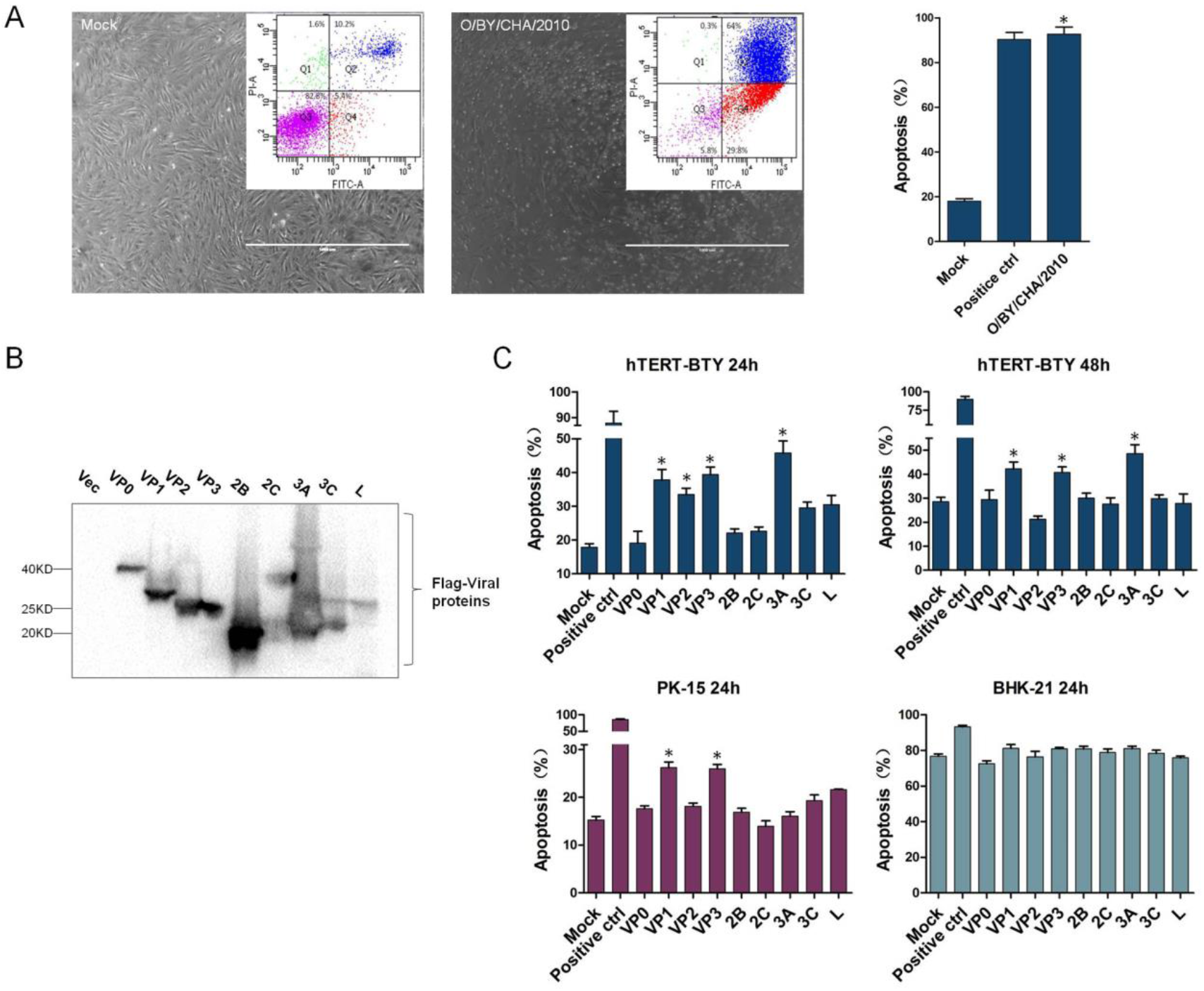
VP1, VP3 and 3A of type O FMDV are inducers of apoptosis. (A) hTERT-BTY cells (5×10^5^ cells/well) were infected or not with O/BY/CHA/2010 at an MOI of 0.05. At 24 h.p.i., the samples were observed under a microscope, and apoptosis was detected by Annexin V-FITC/PI staining and FCM. (B) hTERT-BTY cells (1×10^6^ cells/well) were transfected with FMDV proteins (5μg/well), respectively. At 48 h.p.t, the expression levels of pCAGGS-FMDV-VP0, VP1, VP2, VP3, 2B, 2C, 3A, 3C and L were analyzed by Western blot. (C) hTERT-BTY, PK-15 and BHK-21 cells (5×10^5^cells/well) were transfected with FMDV proteins (3μg/well), respectively. At 24 and 48 h.p.t., respectively, apoptosis was detected by Annexin V-FITC/PI staining and FCM, relative to the negative (empty vector, Mock) and positive (Apoptosis Inducers Kit, Beyotime, China) controls. Experiments were performed in triplicate and repeated three times with similar results. Data are mean ± SD (n=3). Statistical significance was analyzed by Student’s t-test:*P<0.05, **P<0.01; h.p.i., hours post infection; h.p.t., hours post transfection.

### The Gly129 of VP3 random coil is the key apoptotic and autophagic function site

To determine the apoptotic function domain or site of FMDV-VP3, a series of truncation mutants of Flag-FMDV-VP3 expressing plasmids were generated by PCR-based site-directed mutagenesis (Figure 4A). After successful expression of FMDV-VP3 mutants confirmed by Western blot, the above-mentioned plasmids were transfected into hTERT-BTY cells, respectively. The truncated mutants containing the region from carboxyl-terminal 95-to 155-amino acid induced comparable levels of apoptosis, while the other three mutants triggered significantly weaker apoptosis compared with FMDV-VP3 (Figure 4B). The apoptotic function domain in the carboxyl-terminal 106- to 143-amino acid region was subsequently analyzed. Deletion of the region from carboxyl-terminal 124- to 133-amino-acid reduced the apoptotic function of the FMDV-VP3 truncated mutant (VP3 Aa 124-330) compared with the complete VP3 protein (Figure 4C). Then, this region was further interrogated by alanine scanning site-directed mutagenesis. We found that Glycine 129 was essential for FMDV-VP3-induced apoptosis (Figure 4D). To validate its apoptotic function, we mutated this site to Ala in the complete FMDV-VP3 gene and constructed three mutant plasmids (Asia I-VP3 Gly129, A-VP3 Gly129 and O-VP3 Gly129). The results showed that the indicated mutation attenuated the apoptotic function of FMDV-VP3 proteins of type O, Asia I and A FMDV (Figure 4E). Coincidentally, the mutation of Gly129 to Ala remarkably reduced FMDV-VP3 induced LC3-II upregulation. The results indicate that the Gly129 also is a key site of FMDV-VP3 induced autophagy (Figure 4F).

**Figure 4.**
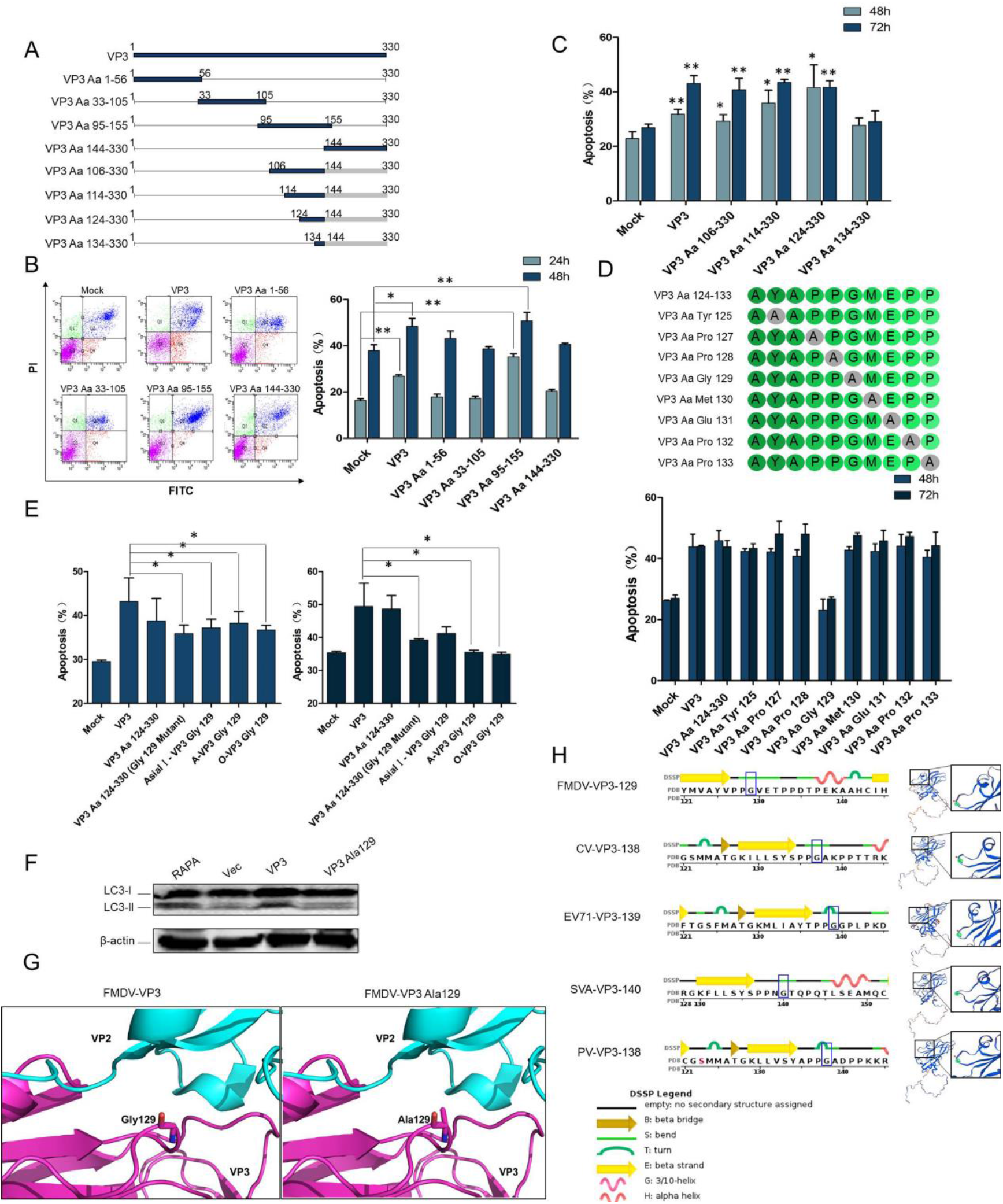
Identification of the apoptotic and autophagic function site of VP3. (A) Schematic representations of a series of Flag-tagged truncated FMDV-VP3 constructs. (B) hTERT-BTY cells (5×10^5^ cells/well) were transfected with FMDV-VP3, indicated FMDV-VP3 mutants and the empty vector, respectively (3μg/well). At 24 and 48 h.p.t., respectively, apoptosis was detected by Annexin V-FITC/PI staining and FCM. (C) hTERT-BTY cells (5×10^5^ cells/well) were transfected with FMDV-VP3, indicated FMDV-VP3 mutants and the empty vector, respectively (3μg/well). At 48 and 72 h.p.t., apoptosis was detected by Annexin V-FITC/PI staining and FCM. (D) hTERT-BTY cells (5×10^5^ cells/well) were transfected with FMDV-VP3, indicated FMDV-VP3 mutants and empty vector, respectively (3μg/well). At 48 and 72 h.p.t., respectively, apoptosis was detected by Annexin V-FITC/PI staining and FCM. (E) hTERT-BTY cells (5×10^5^ cells/well) were transfected with FMDV-VP3, indicated FMDV-VP3 mutants and the empty vector, respectively (3μg/well). At 48 and 72 h.p.t., respectively, apoptosis was detected by Annexin V-FITC/PI staining and FCM. (F) HEK-293T cells (10^5^ cells/well) were transfected with empty vector, FMDV-VP3 or FMDV-VP3 Ala129. At 24 h.p.t., positive control cells were treated with rapamycin (RAPA). The expression levels of caspase-3 were detected by Western blot at 48 h.p.t.. (G) The influences on protein structure of FMDV-VP3 Gly129 mutated to Ala were analyzed by PYMOL software. (H) The location of Gly at FMDV-VP3 (PDB ID:4IV1), CV-VP3 (PDB ID:4Q4V), EV71-VP3 (PDB ID:4RR3), SVA-VP3 (PDB ID:6ADS) and PV-VP3 (PDB ID:5KU0) was analyzed using SWISS-MODEL tool and PDB data bank. Experiments were performed in triplicate and repeated three times with similar results. Data are mean ± SD (n=3). Statistical significance was analyzed by Student’s t-test:*P<0.05, **P<0.01.

The location of Gly129 at picornavirus VP3 was analyzed using SWISS-MODEL tool, PYMOL software and PDB data bank. We find that the Gly129 is located at a bend region of FMDV-VP3 random coil structure. The random coil is easily interacted with amino acids of the surrounding environment, the mutation of Gly129 may change such interaction. In FMDV caspid, for the reason that amino acids side chains get longer, the mutation of Gly129 to Ala has effect on the interaction between VP3 and adjacent VP2 and ultimately remarkable shrunk the volume of viral cavity (Figure 4G). Coincidentally, in the similar location of other picornaviruses, including CV, EV71, SVA and PV, the Gly is conserved (Figure 4H). Above results indicating that the ability of cell death induction may be conserved amongst picornaviruses and this may depend on the specified structure.

In an attempt to investigate the effect of VP3 Gly129 on FMDV-induced apoptosis and its pathological relevance, two recombinant viruses with the apoptotic site of VP3 mutated to Ala or not, namely, rVP3-129/FMDV and rVP3/FMDV, were rescued, respectively. The resultant recombinant viruses were passaged and verified by viral genome sequencing.

### VP3 interacts with p53 and regulates apoptotic and autophagic signaling in a p53 dependent manner

The apoptotic pathway is regulated by a massive and sophisticated network, by which viruses usually manipulate host apoptosis through key apoptotic molecules (40). How FMDV and FMDV-VP3 trigger the apoptotic pathway remains elusive. We firstly examined the contribution of many key apoptotic molecules such as caspases, Bcl-2 related proteins and p53 in FMDV induced apoptosis. The results indicated that caspase proteins, Bcl-2 and p53 mediated signaling pathway might contribute to FMDV induced apoptosis (Figure S2A, S2B). To investigate whether FMDV-VP3 functions by interacting with key apoptotic proteins which directly regulate the apoptotic signaling pathway, hTERT-BTY cells were transfected with the pCAGGS vector/Flag-VP3/Flag-VP3 Ala129. As shown in Figure S2C, endogenous p53 and XIAP were co-precipitated with Flag-FMDV-VP3. However, the Flag-FMDV-VP3 Ala129 failed to interact with p53 and did not affect the interaction between FMDV-VP3 and XIAP. To confirm the interaction between FMDV-VP3 and p53, exogenous and semi-endogenous co-IP were performed. The results of exogenous co-IP showed that only HA-p53 was pulled down by Flag-FMDV-VP3, while no interactions with the Flag-FMDV-VP3 Ala129 and pCAGGS vector were found (Figure 5A). The results of semi-endogenous co-IP showed that under the IgGH band, a 53-KD band corresponding to p53 was evident, whereas no p53 was detected in the empty vector and IgG groups (Figure 5B). Together, these results provided strong evidence that FMDV-VP3 interacted specifically and directly with p53, and the apoptotic site Gly129 was essential for this interaction.

**Figure 5.**
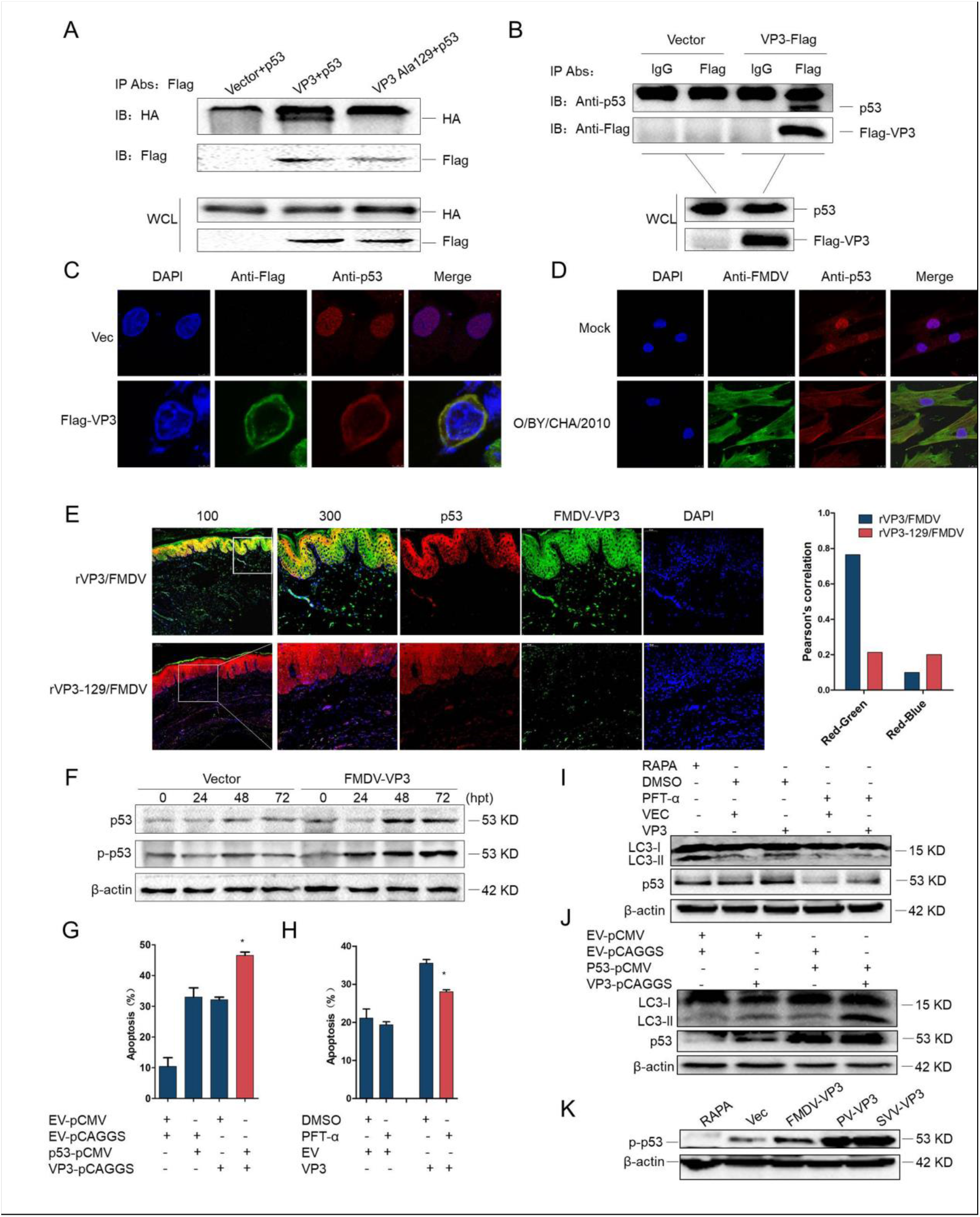
FMDV-VP3 interacts with p53. (A) HEK-293T cells (2×10^6^ cells/well) were co-transfected with 5μg HA-p53 and 5μg empty Flag vector/Flag-FMDV-VP3/Flag-FMDV-VP3-Ala129 for 36 h. Co-immunoprecipitation was performed by using anti-Flag antibody. Anti-p53 antibody was used to detect the precipitated proteins. (B) hTERT-BTY cells (2×10^6^ cells/well) were transfected with 10μg empty Flag vector or Flag-FMDV-VP3 for 48 h, and cell lysates were immunoprecipitated with mouse anti-IgG or anti-Flag antibodies. (C) HEK-293T cells (2×10^5^ cells/well) were co-transfected with 2μg HA-p53 and 2μg Flag-FMDV-VP3/empty Flag vector. At 48 h.p.t., co-localization of HA-p53 (red) and Flag-FMDV-VP3 (green) was observed by confocal microscopy. (D) hTERT-BTY cells (2×10 cells/well) were infected or not with O/BY/CHA/2010 at an MOI of 0.01. At 24 h.p.i., co-localization of p53 (red) and FMDV (green) was observed by confocal microscopy. (E) Guinea pigs were challenged with a high (dilution multiple is 0) dose of the two recovered viruses (200μL/animal). At 7 d.p.c., sections of pathologic tissues were incubated with the indicated antibodies and observed by confocal microscopy. DAPI, blue; FMDV-VP3, green; p53, red. Pearson’s correlation analysis of p53 and FMDV-VP3 or stained nucleus was performed with Image pro plus 6.0. (F) hTERT-BTY cells (5×10 cells/well) were transfected with 3μg empty Flag vector or Flag-FMDV-VP3. At 0, 24, 48 and 72 h.p.t., respectively, the expression levels of p53 and p-p53 were detected by Western blot. (G) hTERT-BTY cells (5×10 cells/well) were transfected with 2μg empty HA vector or HA-p53. At 24 h.p.t., the cells were transfected with 2μg empty Flag vector or Flag-FMDV-VP3 for 48 h. Apoptosis was detected by Annexin V-FITC/PI staining and FCM. (H) hTERT-BTY cells (5×10^5^ cells/well) were pretreated with PFT-α (10μM) or DMSO for 24 h, and transfected with 2μg empty Flag vector or Flag-FMDV-VP3 for 48 h. Apoptosis was detected by Annexin V-FITC/PI staining and FCM. (I) HEK-293T cells (2×10^5^ cells/well) were pretreated with PFT-α (10μM) or DMSO for 24 h, and transfected with 2μg empty Flag vector or Flag-FMDV-VP3 for 48 h, positive control cells were treated with rapamycin (RAPA) for 24h. The expression levels of LC3-I/II and p53 were detected by Western blot at 48 h.p.t.. (J) HEK-293T cells (2×10^5^ cells/well) were transfected with 2μg empty HA vector or HA-p53. At 24 h.p.t., the cells were transfected with 2μg empty Flag vector or Flag-FMDV-VP3 for 24 h. The expression levels of LC3-I/II and p53 were detected by Western blot. (K) HEK-293T cells (10 cells/well) were transfected with empty vector or FMDV-, PV- and SVA-VP3. At 48 h.p.t., the expression levels of p-p53 were detected by Western blot. Except for animal experiments, all assays were performed in triplicate and repeated three times with similar results. Data are mean ± SD (n=3). Statistical significance was analyzed by Student’s t-test:*P<0.05, **P<0.01. EV, Empty Vector.

To further visualize the interaction between FMDV-VP3 and p53, immunofluorescence was carried out. As illustrated in Figure 5C, stabilized p53 accumulated in the nucleus in vector infected HEK293T cells, whereas upon expression of FMDV-VP3, p53 was translocated from the nucleus to the cytoplasm and co-localized with FMDV-VP3. The FMDV also co-localized with endogenous p53 in the cytoplasm (Figure 5D). In in vivo experiments, FMDV-VP3 was co-localized with endogenous p53 in the cytoplasm of rVP3/FMDV-challenged guinea pig tissues (Figure 5E). However, in most tissue cells of rVP3-129/FMDV-challenged guinea pigs, p53 mainly accumulated in the nucleus and to a lesser extent, co-localized with FMDV-VP3 in the cytoplasm. These results revealed that FMDV-VP3 contributes to p53 translocation from the nucleus into the cytoplasm, and Gly129 is essential for the co-localization of FMDV-VP3 and p53. A similar phenomenon was observed in both cells and animals infected by the FMDV.

Then, a VP3-induced increase in p53 protein levels was detected in FMDV-VP3 but not empty vector transfected hTERT-BTY cells (Figure 5F). We next explored which post-translational modification types FMDV-VP3 employed to facilitate the activation and accumulation of p53. hTERT-BTY cells were transfected with FMDV-VP3 plasmids or the empty vector, and the results showed that FMDV-VP3 promoted p53 phosphorylation as transfection time increased. Furthermore, overexpression of p53 significantly enhanced FMDV-VP3-induced apoptosis in hTERT-BTY cells (Figure 5G), suggesting that p53 is crucial for VP3-induced apoptosis. Moreover, the apoptosis level of FMDV-VP3-transfected cells pre-treated with PFT-α was significantly lower than that of the DMSO group (Figure 5H). The results indicated that p53 directly promoted FMDV-VP3-induced apoptosis. The effect of p53 on FMDV-VP3-induced autophagy was also investigated. The results of Western blot indicated that the LC3-II upregulation induced by FMDV-VP3 was substantially declined with p53 pathway inhibition (Figure 5I) while that was further enhanced by p53 overexpression (Figure 5J). For other picornaviruses, containing PV and SVV, were proved to upregulated the expression of phosphorylated p53 (Figure 5K). All the data indicated that p53 is a crucial node in VP3-induced apoptosis and autophagy pathway, and the function may be conserved amongst picornaviruses.

### FMDV-VP3 promotes p53 interaction with the pro-apoptotic protein Bad, and triggers the mitochondrial apoptotic pathway

Previous studies suggested that p53 translocates from the nucleus into to cytoplasm (41). Under many pro-apoptotic conditions, p53 is transferred to the mitochondria and directly interacts with critical Bcl-2 related proteins such as Bax, Bak, Bcl-XL, Bcl-2 and Bad (42, 43). To assess the downstream events following the interaction between FMDV-VP3 with p53, a series of co-immunoprecipitation experiments were performed to screen for downstream proteins. hTERT-BTY cells were co-transfected with the Flag-FMDV-VP3-expressing plasmid or p53-HA/pCMV vector. The results showed that only Bad was pulled down by p53-HA, but not with the pCMV-HA vector (Figure 6A).

**Figure 6.**
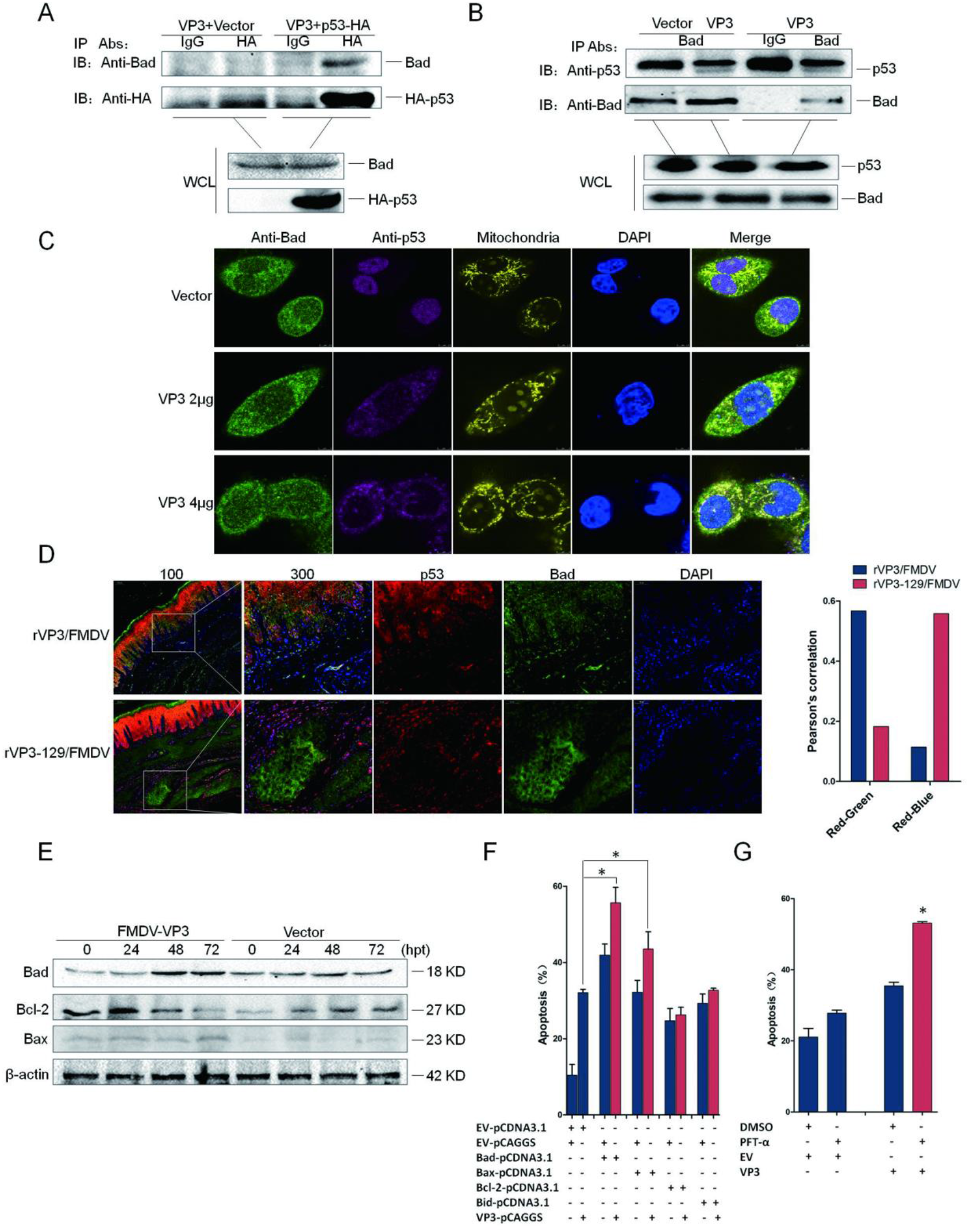
p53 interacts with Bad in the presence of FMDV-VP3. (A) hTERT-BTY cells (2×10^6^ cells/well) were co-transfected with Flag-FMDV-VP3 (5μg) and HA-p53/empty HA vector (5μg) for 48 h. The lysates were immunoprecipitated with mouse anti-IgG or anti-HA antibody. The eluted fractions were detected with anti-Bad antibody. (B) hTERT-BTY cells (2×10^6^ cells/well) were transfected with empty Flag vector or Flag-FMDV-VP3 (10μg). At 24 h.p.t., cell lysates were immunoprecipitated with anti-Bad or anti-IgG antibody. The precipitated proteins were blotted with anti-p53 antibody. (C) HEK-293T cells (10 cells/well) were transfected with empty Flag vector or increasing quantities of Flag-FMDV-VP3 (2μg or 4μg). At 24 h.p.t., cell mitochondria were labeled with the mitochondrial probe (500 nM) for 40 min. Then, cells were fixed and incubated with anti-endogenous p53 and Bad antibodies, respectively. Co-localization of p53 (purple), Bad (green) and the mitochondria (orange) was observed by confocal microscopy. (D) Guinea pigs were challenged with a high (dilution multiple is 0) dose of the two recovered viruses (200μL/animal). At 7 d.p.c., the lesions were incubated with the indicated antibodies and examined by confocal microscopy. DAPI, blue; Bad, green; p53, red. Pearson’s correlation analysis of p53 and Bad or stained nucleus was analyzed with Image pro plus 6.0. (E) hTERT-BTY cells (5×10 cells/well) were transfected with 3μg empty Flag vector or Flag-FMDV-VP3. At 0, 24, 48 and 72 h.p.t., respectively, the expression levels of Bad, Bcl-2 and Bax were detected by Western blot. (F) hTERT-BTY cells (5×10 cells/well) were transfected with 2μg empty vector and Bad, Bax, Bcl-2, and Bid-expressing plasmids, respectively. At 24 h.p.t., the cells were transfected with 2μg empty Flag vector or Flag-FMDV-VP3 for 48 h. Apoptosis was detected by Annexin V-FITC/PI staining and FCM. (G) hTERT-BTY cells (5×10^5^ cells/well) were pretreated or not with TW-37 (10μM) for 24 h, and transfected with 2μg empty Flag vector or Flag-FMDV-VP3 for 48 h. Apoptosis was detected as above. Except for animal experiments, assays were performed in triplicate and repeated three times with similar results. Data are mean ± SD (n=3). Statistical significance was analyzed by Student’s t-test: *P<0.05, **P<0.01.

To assess whether VP3 expression is essential for the interaction between p53 and Bad, hTERT-BTY cells were transfected with Flag-FMDV-VP3 or the vector. As shown in Figure 6B, p53 was pulled down by Bad in the presence of FMDV-VP3; nevertheless, there was no interaction between p53 and Bad in the vector-transfection group. Moreover, to examine whether VP3 enables Bad to co-localize with p53 in the mitochondria, the Flag-FMDV-VP3-expressing plasmid or vector was transfected into HEK293T cells. Bad is normally scattered in the cytoplasm, whereas p53 is localized in the nucleus. However, upon expression of FMDV-VP3, p53 was translocated from the nucleus to the cytoplasm, and co-localization of p53 and Bad in the mitochondria was observed (Figure 6C). In in vivo experiments, footpad lesions were used to map the distribution of p53 and Bad. Double immunofluorescent staining showed that both endogenous p53 and Bad co-localized in the cytoplasm after challenge with rVP3/FMDV. In contrast, Bad in tissue cells of rVP3-129/FMDV-challenged animals displayed diffused distribution, and co-localization with p53 was merely detected (Figure 6D).

The anti-apoptotic protein Bcl-2 and pro-apoptotic proteins Bad and Bax are key molecules responsible for the intrinsic mitochondria apoptotic pathway. Experiments were performed to determine the importance of Bad, Bcl-2 and Bax in FMDV-VP3-induced apoptosis. The expression levels of Bad and its downstream protein Bax were increased, while Bcl-2 amounts were decreased by FMDV-VP3 transfection (Figure 6E). Next, we found that FMDV-VP3 notably increased apoptotic rates in Bad and Bax overexpressing cells. However, there was no change of FMDV-VP3-induced apoptosis in cells overexpressing another Bcl-2 related protein, Bid (Figure 6F). Then, the effect of Bcl-2 pathway inhibition on FMDV-VP3-induced apoptosis was investigated. The results indicated that apoptosis induced by FMDV-VP3 was substantially increased in cells with inhibited Bcl-2 pathway (Figure 6G). Collectively, these results indicated that FMDV-VP3 induces apoptosis via the mitochondrial pathway mediated by Bad, Bax and Bcl-2.

The downstream components of the complex comprising p53 and Bad in the FMDV-VP3-dependent apoptosis pathway were then determined. The results revealed that the levels of Fas, Akt and cleaved caspases −3, 8 and 9 were significantly increased by FMDV-VP3 rather than the empty vector, as well as cytoplasmic cyt-c amounts, whereas mitochondrial cyt-c levels were notably reduced (Figure S3A). Since many key proteins of the intrinsic and extrinsic apoptotic pathways were regulated by FMDV-VP3, to examine which of the aforementioned proteins are downstream components of p53 in the FMDV-VP3-dependent apoptosis pathway, hTERT-BTY cells were pre-treated with the p53 specific inhibitor PFT-α for 24 h and transfected with FMDV-VP3 or empty vector. Compared with cells untreated with PFT-α, FMDV-VP3-induced upregulation or downregulation of Bad, Bax, Bcl-2, cytoplasmic cyto-c and cleaved caspase-3 was significantly suppressed in the PFT-α pretreatment group, but p53 inhibition had no effect on FMDV-VP3-induced upregulation of Akt (Figure S3B). This implies that Bad, Bax, Bcl-2, cytoplasmic cyto-c and caspase-3 are downstream molecules of p53 in the FMDV-VP3-dependent apoptosis pathway, where FMDV-VP3 might activate another apoptotic pathway mediated by Akt.

### Gly129 of VP3 is essential for VP3-induced apoptosis and autophagy that enhance FMDV pathogenicity

To determine the effect of the VP3 Gly129 residue on FMDV internalization, the two recombinant viruses were incubated with hTERT-BTY and PK-15 cells for 1 h. Unabsorbed and surface-bound viruses were removed, and rRT-PCR was used to quantitate the internalized FMDV. The results showed no significant influence on viral internalization in both hTERT-BTY and PK-15 cells with VP3 Gly129 mutated into Ala (Figure 7A). On the premise of similar internalization, hTERT-BTY cells were infected with the aforementioned viruses to analyze the impact of the VP3 Gly129 residue on FMDV-induced cellular apoptosis. Our data showed that the apoptosis rate of rVP3-129/FMDV infected cells was drastically lower than that of the rVP3/FMDV infected group (Figure 7B). This was further corroborated by TUNEL assays on major pathologically-changed tissues, in which the percentages of TUNEL positive cells in both high (0) and low (−5) dose groups challenged by rVP3-129/FMDV were lower than those of the corresponding dose groups challenged by rVP3/FMDV (Figure 7C). Consistently, results of immunofluorescent staining on footpad lesions showed that the positive cells percentage and mean fluorescence intensity of FMDV, p53, cleaved-caspase3 and LC3-I/II are remarkably higher in rVP3/FMDV challenged groups than the corresponding dose groups challenged by rVP3-129/FMDV. It is important to emphasize that the statistical gap is particularly noticeable in middle and later stage (15 d.p.c.) of FMDV infection (Figure 7D). The mutation of Gly129 to Alu remarkably reduced rVP3/FMDV induced LC3-II upregulation was further corroborated by Western blot (Figure 7E). Taken together, the decreased ability to induce apoptosis and autophagy by FMDV in vivo was directly attributable to the mutation of the VP3 Gly129.

**Figure 7.**
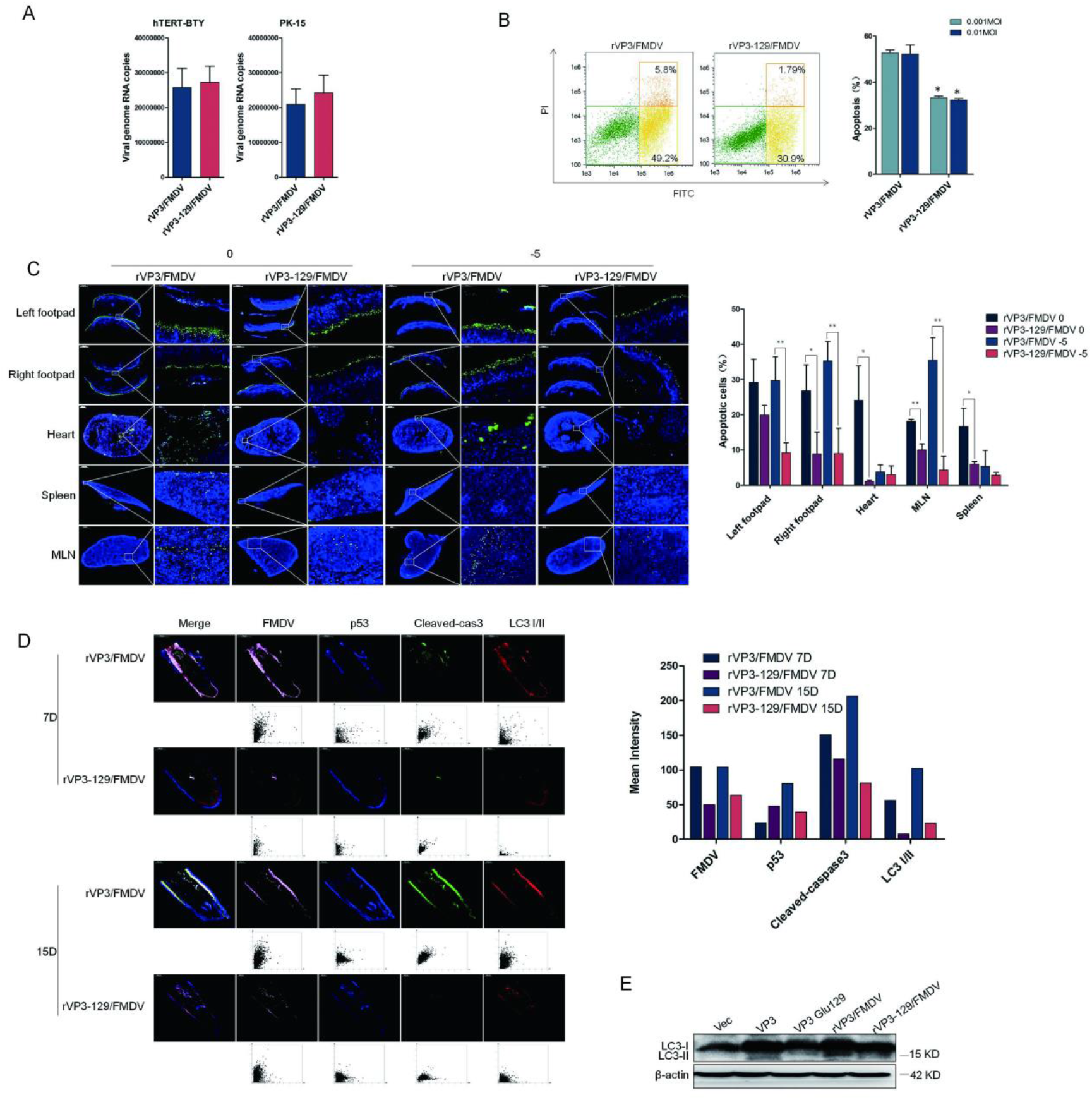
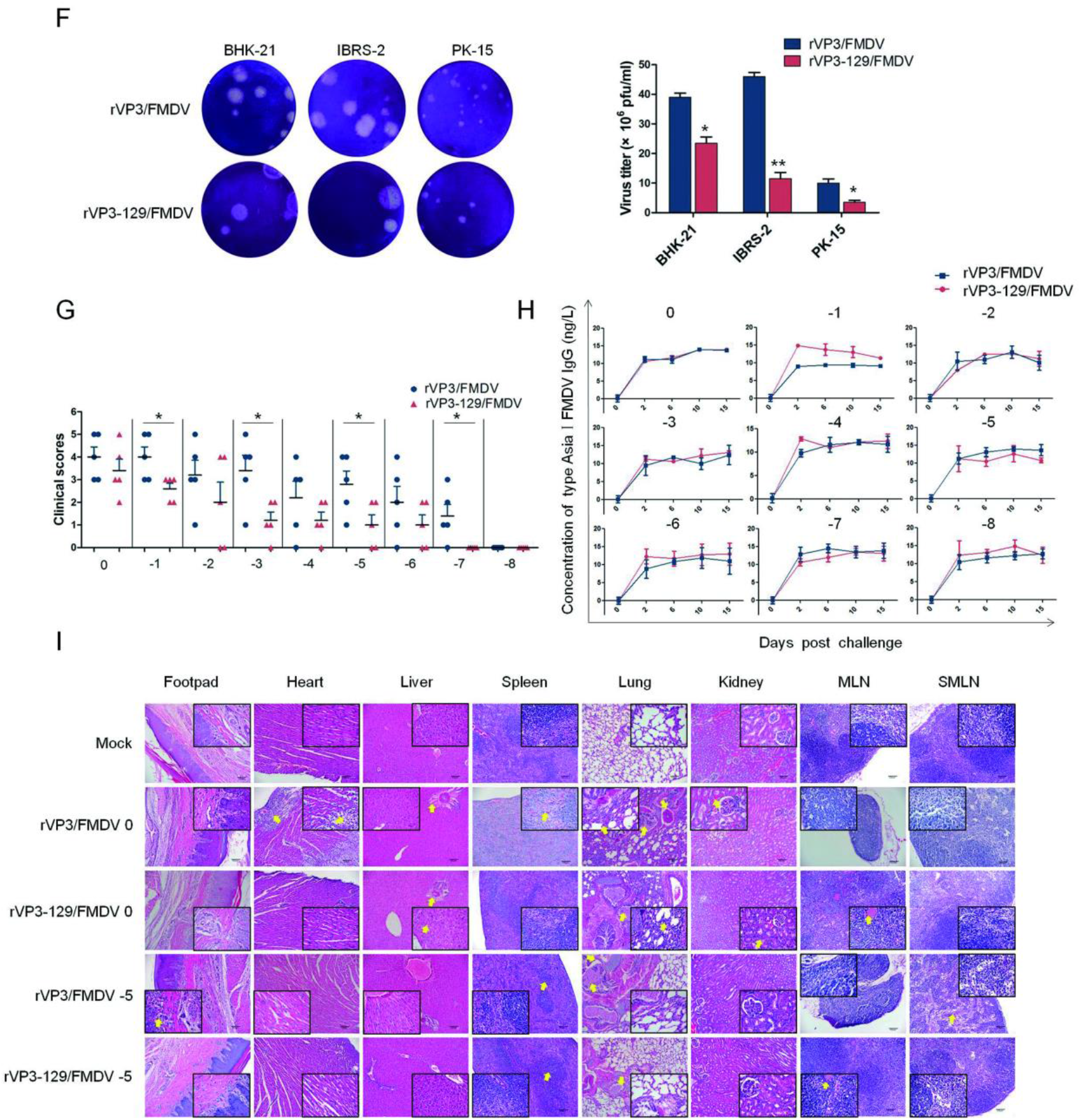
The apoptotic site of VP3 is essential for FMDV apoptotic and autophagic function, titer and pathogenicity. (A) hTERT-BTY and PK-15 cells (5×10 cells/well) were infected with the two rescued viruses at a dose of 1 ×108 viral genomic RNA copies for 1 h at 37°C, and unabsorbed viruses were washed with ice-cold Hanks solution three times. To remove the surface-bound viruses, the cells were digested with trypsin-EDTA (GIBCO) for 5 min. Then, the cells were washed three times with Hanks balanced salt solution and rRT-PCR was used to quantitate the internalized FMDV. (B) hTERT-BTY cells (5×10^5^ cells/well) were infected with the two rescued viruses at an MOI of 0.001 or 0.01, at 24 h.p.i., and apoptosis was detected by Annexin V-FITC/PI staining and FCM. (C) Guinea pigs were challenged with high (dilution multiple is 0) and low (dilution multiple is 10) doses of the two recovered viruses (200μL/guinea pig), respectively, and apoptotic rates of major pathologically-changed tissues, including the left and right footpads, heart, spleen and MLN, at 7 d.p.c. were analyzed by the TUNEL assay. The percentage of apoptotic cells was analyzed with Image pro plus 6.0. (D) Guinea pigs were challenged with high (dilution multiple is 0) doses of the two recovered viruses (200μL/guinea pig), respectively, At 7 and 15 d.p.c., the footpad lesions were incubated with the indicated antibodies and examined by confocal microscopy. FMDV, pink; p53, blue; cleaved-caspase3, green; LC3-I/II, red. The positive cells were analyzied respectively by HISTOQUEST soteware. (E) HEK-293T cells (10^5^ cells/well) were transfected with empty vector, FMDV-VP3, FMDV-VP3 Ala129 or infected with the two recovered viruses at an MOI of 0.01. The expression levels of LC3-I/II were detected by Western blot at 48 h.p.t. or 36 h.p.i.. (F) Cell monolayers were infected with the two rescued viruses at 37°C for 1 h, and 0.6% gum Tragacanth overlay was added. After 48 h.p.i., the cells were stained with crystal violet. Plaques were captured, and PFUs were counted. (G) Guinea pigs were challenged with the indicated doses of the two recovered viruses (200μL/guinea pig). Clinical symptoms were scored using the double-blind method at 3 d.p.c. (H) FMDV-specific antibody titers of the challenged guinea pigs at 0, 3, 6, 10 and 15 d.p.c. were measured with a guinea pig FMDV IgG ELISA kit. (I) Guinea pigs were challenged with high (dilution multiple is 0) and low (dilution multiple is 105) doses of the two recovered viruses (200μL/guinea pig); the Mock group was injected with PBS. Histology of the heart, liver, spleen, lung, kidney, MLN, submaxillary lymph nodes (SMLN) and pathological tissues of the footpads at 7 d.p.c. were analyzed by H&E staining (100×, 400×magnification). Except for animal experiments, assays were performed in triplicate and repeated three times with similar results. Data are mean ± SD (n=3). Statistical significance was analyzed by Student’s t-test:*P<0.05, **P<0.01; d.p.c., days post challenge.

We next performed plaque assays in BHK-21, IBRS-2 and PK-15 cells. The results indicated that the mutation of the apoptotic site substantially reduced FMDV titers (PFU/mL) in all tested cell lines (Figure 7F).

In in vivo experiments, clinical signs of challenged guinea pigs were observed daily. The clinical scores of the −1, −3, −5 and −7 challenge groups administered rVP3-129/FMDV were significantly lower than those of the corresponding rVP3/FMDV groups (Figure 7G), indicating that VP3 Gly129 mutation directly reduces the virulence and pathogenicity of the FMDV.

FMDV-specific antibody titers were measured. There was no significant difference in antibody levels between comparable dose challenge groups, with the exception of the −1 group, where the IgG levels of the rVP3-129/FMDV-challenge group were noticeably higher than those of rVP3/FMDV-challenged guinea pigs (Figure 7H).

To assess the histopathology of animals challenged with the two FMDV strains, H&E staining of major tissues was performed. In the rVP3/FMDV 0-challenge group (Figure 7I), pathological changes occurred in almost all major organs of guinea pigs, e.g. extensive necrosis, degeneration, and inflammatory cell infiltration. On the contrary, the extent of tissue damage was markedly reduced in rVP3-129/FMDV 0-challenged guinea pigs, where the heart tissues appeared normal. Minor pathological changes were observed in the liver, lung, kidney, mesenteric lymph nodes (MLN) and claw lesions. Likewise, the extent of tissue damage of low dose rVP3-129/FMDV-5-challenged guinea pigs was also significantly reduced compared with that of the rVP3/FMDV-5-challenge group. Taken together, these results indicated that the mutation of the VP3 apoptotic site could be an important determinant of reduced FMDV pathogenicity.

### VP3-induced apoptosis occurs at the middle and late stages of infection, and promotes FMDV replication

Apoptosis induced by viruses may either stimulate or inhibit viral replication (44). Viral growth curves in FMDV permissive animal cell lines such as hTERT-BTY and PK-15 cells were generated to compare the propagation capacities of the two recombinant viruses. As shown in Figure 8A, rVP3-129/FMDV exhibited a diminished ability to replicate in hTERT-BTY and PK-15 cells compared with rVP3/FMDV, suggesting that VP3 Gly129 is directly related to viral proliferation capacity.

**Figure 8.**
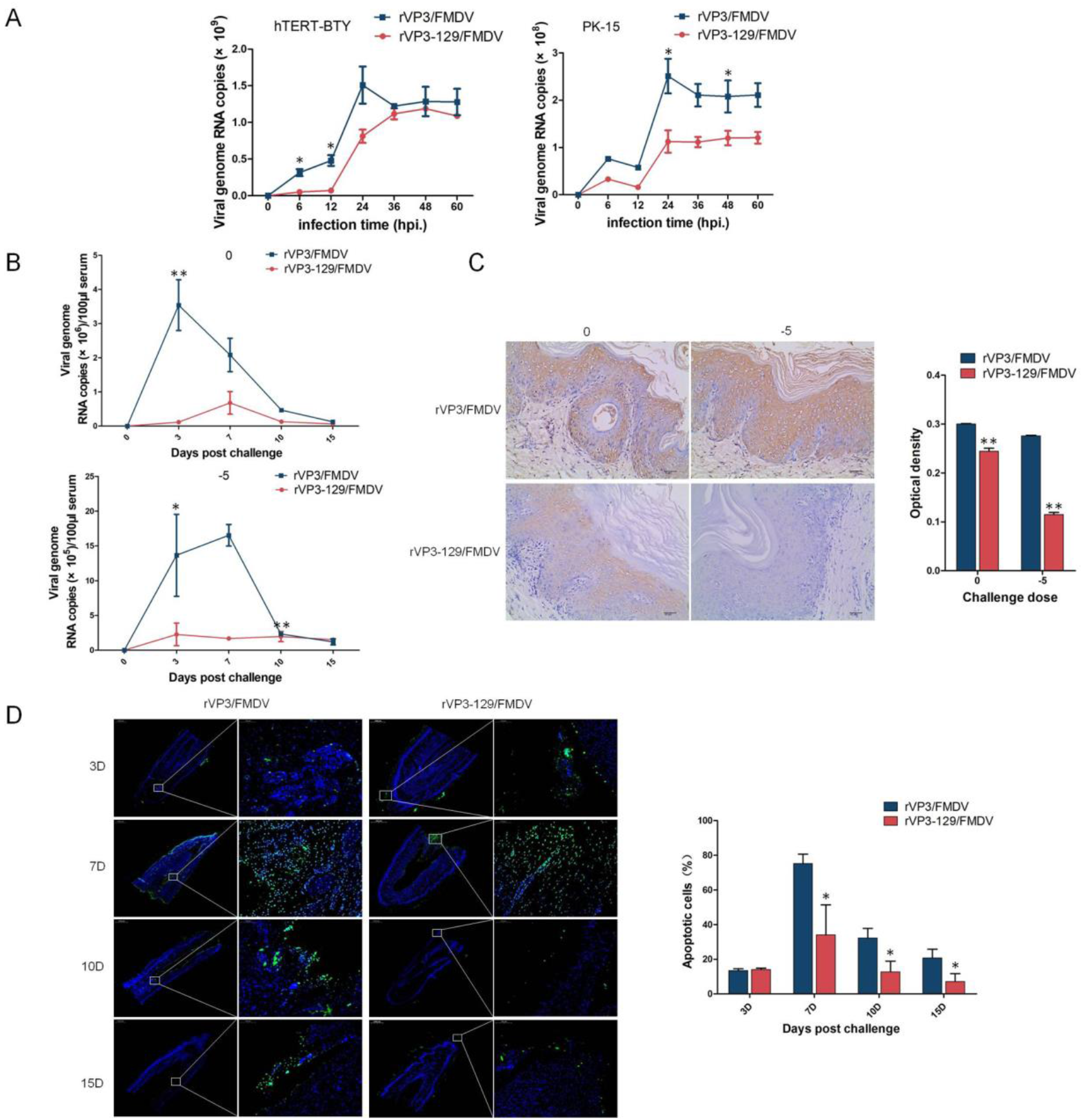
Apoptosis and autophagy induced by VP3 play a critical role in FMDV spread and replication. (A) hTERT-BTY and PK-15 cells (5×10 cells/well) were infected with the two rescued viruses at a dose of 1×106 viral genomic RNA copies for 1 h, and unabsorbed viruses were washed. Cells and supernatants were harvested at 0, 6, 12, 24, 36, 48 and 60 h.p.i., respectively, and viral RNA amounts were determined by RT-PCR. Experiments were performed in triplicate and repeated three times with similar results. Date are mean ± SD (n=3). (B) Guinea pigs were challenged with high (dilution multiple is 0) and low (dilution multiple is 105) doses of the two recovered viruses (200μL/guinea pig). Viral RNA copies in serum were detected by RT-PCR. (C) Guinea pigs were challenged with high (dilution multiple is 0) and low (dilution multiple is 105) doses of the two recovered viruses (200μL/guinea pig). FMDV of pathogenic claw tissues were determined by IHC at 7 d.p.c. Statistical analysis of FMDV amounts was performed based on IHC staining results in 3 randomly fields per section. (D) Guinea pigs were challenged with the two recovered viruses (dilution multiple is 10^4^), and apoptotic rates of footpad lesions at 3, 7, 10 and 15 d.p.c., respectively, were analyzed by the TUNEL assay. Statistical significance was analyzed by Student’s t-test:*P<0.05, **P<0.01.

FMDV replication levels in blood were also assessed. The viremia levels of groups challenged by the two rescued viruses increased over time and peaked at 6 days post challenge (d.p.c.). Compared with rVP3/FMDV-challenged guinea pigs, the rVP3-129/FMDV-challenge group showed significantly lower viremia levels (Figure 8B). In addition, FMDV amounts in footpad lesions in both challenge groups were analyzed by IHC. The data indicated that the viral load in rVP3-129/FMDV-challenged guinea pigs was substantially lower than in rVP3/FMDV-challenged animals (Figure 8C).

Next, we assessed the rates of apoptosis over time in lesion sites of guinea pigs challenged by the two FMDV strains. There was no significant difference in apoptotic rate between the two groups at the early stage (3 d.p.c.). In contrast, at the mid and late stages (7–15 d.p.c.), apoptotic rates of rVP3-129/FMDV-challenge groups were substantially lower than those of rVP3/FMDV-challenged animals (Figure 8D).

**Figure 9.**
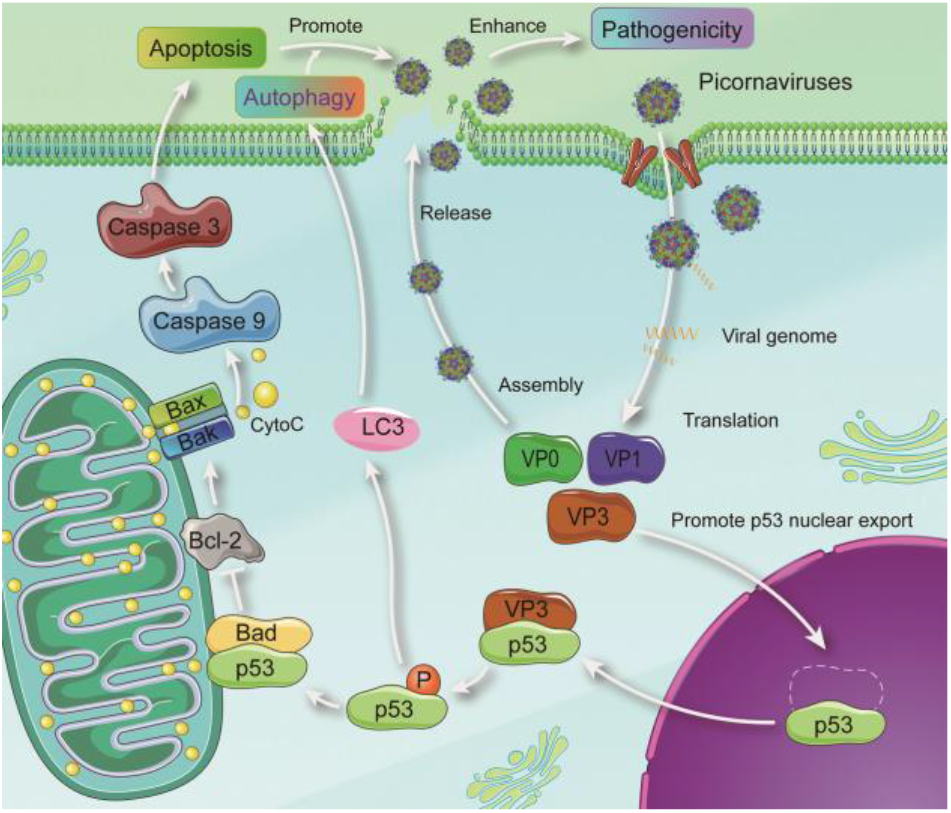
Hypothetical schematic model of the mechanism of VP3-induced cell death. As a stress signal, picornaviruses infection promotes p53 translocation from the nucleus into the cytoplasm and interacts with VP3. Activated p53 subsequently translocates to the mitochondria and interacts with Bad, which triggers the Bcl-2 family dependent intrinsic apoptotic signaling pathway. Meanwhile, activated p53 triggers the LC3 dependent autophagy. Finally, apoptosis and autophagy facilitate viral replication and enhance pathogenicity.

## DISCUSSION

Most viruses have the capability to modulate apoptosis and autophagy in the host for their replication and dissemination (45, 46). Picornaviruses caused a variety of severe diseases in humans and livestock. To promote replication, the viral proteases 2A, 3C and 3CD of picornaviruses cleave proapoptotic proteins or nucleoporins to modulate apoptosis; meanwhile, 2B protein alters intracellular ion signaling and controls apoptosis (47–49). Viral infection also is tightly related to autophagy. Poliovirus proteins 2BC increases LC3 lipidation and 3A inhibits autophagosome movement along microtubules (25, 50). CVB induces autophagy to promote its replication (51). FMDV protein VP2 induces autophagy to enhance replication (31). However, whether other picornaviruses proteins regulate cell death and the association of cell death with viral pathogenicity remains unclear. In the present study, to explore novel cell death regulation proteins of picornaviruses, bioinformatics prediction approaches were performed. The functions of proteins to regulate cell death are often related to its hydrophobic regions. BH3-only proteins interact with BAK at the canonical hydrophobic groove (52). Bcl-xL interacts with other apoptosis regulation proteins through a large, primarily hydrophobic binding groove. Phosphorylation of the hydrophobic motif ser 662 is an essential step in the activation of the cell survival and death regulation protein protein kinase C delta (PKCd) (53). The transmembrane hydrophobic sequences of CVB non-structure protein 2B is essential for the induction of autophagy (54). In view of this, we firstly predict the hydrophobic property of FMDV, PV and SVA proteins. VP3 proteins of the three virus were selected as the main focus because which not only have high hydropathicity score, but also have more amino acids with the top score. The VP3 of FMDV, PV and SVA has similar structure and they can induce both apoptosisand autophagy by verification.

FMDV was chosen as a model to understand further molecular mechanism of VP3 inducted cell death. FMDV is cytocidal in vivo and in vitro (55), but the main type of FMDV-induced cell death is still unclear. Here we found that FMDV can induce both apoptosis and autophagy *in vivo*, but the main type of cell death is apoptosis. *In vitro*, VP3 of FMDV, PV and SVA was identified as inducer of apoptosis and autophagy. VP3, a component of picornavirus capsid, makes a substantial contribution to capsid stability and contains receptor recognition sites and important neutralizing epitopes (56). Which is also closely related to virulence and innate immune response. C-terminal aa 111-220 of FMDV-VP3 are essential for VP3 interaction with VISA to inhibit innate immune response (57). The His124 of serotype A FMDV-VP3 mutated to Asp increases viral acid resistance (58). The Arg56 of FMDV-VP3 is closely associated with virulence (59). The aa 59, 62, 67 and 176-190 of EV-71-VP3 are conformational neutralizing epitopes (60, 61). The function of other aa of VP3 is not clear. We found that FMDV-VP3 significantly induces apoptosis and autophagy, and the Gly129 locates at a bend region of random coil structure is the apoptotic and autophagic function site. The mutation of Gly129 may change the interaction between VP3 and its close amino acids around which. More importantly, the Gly is conserved at the similar location of other picornaviruses, including CV, EV71, SVA and PV, indicating that the Gly of picornaviruses VP3 random coil has similar function that dependent on the special structure location.

Viruses usually modulate a key molecule to manipulate host cell death (40). How FMDV-VP3 triggers the apoptotic and autophagic pathway remains unclear. In the present study, VP3 specifically interacted with p53 in the cytoplasm both *in vivo* and *in vitro*. Intriguingly, the interaction and co-localization disappeared when Gly 129 was mutated. The tumor suppressor p53 is considered a key molecule in the regulation of apoptosis, autophagy and growth arrest in the cellular response to replicative and environmental stresses. (41, 62). The multifunction of p53 depending of its intracellular localization. Indeed, p53 is a well-characterized transcription factor that transactivates a number of genes related to apoptosis and autophagy, such as Bid (63), Bax (64), Fas (65), Noxa (66), Puma (67), damage-regulated autophagy modulator (Dram) (68), PTEN-induced kinase 1 (PINK1)(69) and *Isg20L1*(70). Indisputably, the sub-cellular location in which p53 performs the above function is the nucleus. However, in addition to its nuclear activity, it has also been reported that p53 directly interacts with viral proteins to trigger the p53-independent cell death pathway under certain circumstances. For instance, p53 interacts with the U14 protein of human herpesvirus and plays an important role in viral infection (71). The interaction between p53 and hepatitis B virus X protein results in the abrogation of apoptosis (72). The BM2 protein of influenza B virus interacts with p53 and blocks p53-mediated apoptosis (73). In addition, previous studies suggested that p53 can translocate from the nucleus to the cytoplasm (41, 74). Here we demonstrated that VP3 contributes to p53 translocation from the nucleus into the cytoplasm. Challenge with rVP3/FMDV led to p53 co-localization with FMDV-VP3 in the cytoplasm of tissue cells. However, in the absence of the Gly129, p53 was predominantly localized to the nucleus, with no apparent co-localization between p53 and FMDV-VP3 in the tissues of rVP3-129/FMDV challenged guinea pigs.

We also found that p53 levels were increased in case of VP3 expression, and phosphorylation of p53 was promoted by VP3. More importantly, VP3-induced apoptosis and autophagy were significantly enhanced by p53 overexpression, but decreased by the p53 specific inhibitor PFT-α. In response to stress, p53 undergoes post-translational phosphorylation that is believed to regulate its accumulation and activation, and accumulation is prominently linked to mitochondrial translocation of p53 (75, 76). Under many stress conditions, p53 translocates to the mitochondria and directly interacts with critical Bcl2 related proteins, which regulate both apoptosis and autophagy, such as Bax, Bak, Bcl-XL, Bcl2 and Bad (42, 43, 77). Meanwhile, p53 has been suggested to function like a BH3-only protein in the mitochondria and induce mitochondrial outer membrane permeabilization, leading to the release of mitochondrial cytochrome c (78). As shown above, Bad was pulled down by p53 in the presence of VP3, which enables Bad to co-localize with p53 in the mitochondria both in HEK-293T cells and tissues of rVP3/FMDV challenged guinea pigs. Mutation of Gly 129 greatly reduced the co-localization level of p53 and Bad. These results suggested that wild-type VP3 may act as an inducer of p53 translocation to the mitochondria and the interaction with Bad. In addition, we also validated that VP3 can upregulate Bad and ultimately resulting in apoptotic and autophagic cell death of host cells.

Apoptosis, autophagy and unfolded protein response are the main mechanisms involved in viral pathogenesis (44). In many viruses such as yellow head virus (79), mouse hepatitis virus (80) and Japanese encephalitis virus (81), virus induced apoptosis is considered a virulence factor and may promote viral pathogenicity and mortality (82–85). Influenza A virus induced apoptosis is a cause of organ damage (86). H5N1- or H1N1-induced distal lung epithelial cell apoptosis is a classical feature of acute respiratory disease syndrome (58). HIV-1-induced autophagy in different stages may stimulate or inhibit viral infection and pathogenesis (87–89). Whether VP3 of picornaviruses induced apoptosis and autophagy are relevant to biological characteristics of the virus remains unclear. Here, two recombinant viruses with VP3 Gly129 mutated to Ala or not termed rVP3-129/FMDV and rVP3/FMDV, respectively, were rescued. In *in vitro* experiments, mutation of Gly129 significantly reduced the viral titer, apoptosis and autophagy levels. In *in vivo* experiments, guinea pigs, as a FMDV susceptible animal model, were challenged by the two recombinant viruses, and mutation of Gly129 directly contributed to diminished ability of the FMDV to induce apoptosis and autophagy in organs. However, whether the degree of VP3-induced apoptosis and autophagy are positively associated with organ damage remains unclear. The present results indicated that Gly129 mutation does not affect the internalization process of the FMDV. Therefore, VP3 Gly129 mutation significantly relieved the clinical symptoms of the FMDV, while FMDV-specific antibody titers were negligibly affected. Furthermore, this mutation effectively alleviated the degree of damage in multiple organs. Taken together, these findings indicate that FMDV VP3-induced apoptosis and autophagy played crucial roles in FMDV virulence and pathogenicity.

In general, viral infection triggered apoptotic and autophagic cell death are regulated by the host’s active defense system. The self-destructive apoptosis may cause abortion of unassembled virus. Since apoptotic cell death impedes viral replication, viruses, to maximize viral propagation, express anti-apoptotic proteins to delay or block apoptosis. This process makes more time for viral assembly and replication before the death of host cells (2, 90). Such apoptosis usually occurs at the early stage of infection and reduces viral replication (91–94). Moreover, a number of viruses that have not evolved anti-apoptotic or evasion mechanisms, including influenza virus, may encourage apoptosis to promote viral replication (95, 96). For example, Chikungunya virus triggers apoptosis and utilizes the resulting apoptotic blebs to circumvent host cell defense mechanisms, thereby facilitating viral dissemination and replication (97). Autophagy also acts as both anti-viral and pro-viral roles in viral infection. As a cell-intrinsic defensive mechanism, host cell may degrade viral components by autophagy pathway(98, 99). As a cellular survival mechanism, autophagy restrains the spread of virus from the primary infection site to adjacent uninfected cells (5, 98). For RNA virus, autophagosome as cellular membranous structure, generally serve as a platform for membrane-associated replication factories to replicate and assemble(6). In the present study, mutation of Gly 129 significantly reduced not only FMDV replication *in vivo* and *in vitro*, but also the viral load of the FMDV in pathogenic claw tissues. These results indicated that VP3-induced apoptosis and autophagy played critical role in promoting viral proliferation. As previously reported, many viruses use apoptosis to kill cells at the late stage of infection. During the process, progeny virions are encapsulated into apoptosis bodies and autophagic vesicles that rapidly infect the surrounding cells (100). In this manner, the virus can spread but could not be destroyed by virus-induced host’s inflammatory response, immune response and protease digestion (2, 40). For instance, in order to promote viral spread, the Adenovirus E4 or F4 protein can kill cells at the end of the infectious cycle (101). Here we observed that Gly 129 mutation significantly reduced the apoptotic and autophagic capacity of the FMDV at the mid and late stages rather than the early stage of infection, suggesting that FMDV-VP3 induced apoptosis and autophagy occurs at the mid and late stages of infection, which could be a major approach for promoting the spread of viral progeny and infection of neighboring cells, thereby enhancing viral replication.

In summary, the FMDV VP3 protein directly interacts with p53 during infection, and promotes p53 translocation to the mitochondria and its interaction with Bad. This then triggers Bcl-2 family-dependent intrinsic apoptosis and LC3-dependent autophagy signaling pathway. The apoptosis and autophagy occurred at the mid and late stages of FMDV infection, thereby enhancing viral replication and pathogenicity. The apoptotic and autophagic function may be conserved amongst picornaviruses as the functional site Gly is conserved at the similarly location of other picornaviruses.

## AVAILABILITY

All relevant data are available from the corresponding authors upon request.

## ACKNOWLEDGMENTS

We thank the Facility Center Department, Lanzhou Veterrinary Research Institute and Analysis and Test Group, Center for Technical Development and Analysis Service, Institute of Modern Physics, for helpful support.

